# High-resolution Profiling of Bacterial and Fungal Communities using Pangenome-Informed Taxon-Specific Long-Read Amplicons

**DOI:** 10.1101/2023.07.17.549274

**Authors:** Luzia Stalder, Monika Maurhofer, Daniel Croll

**Affiliations:** Laboratory of Evolutionary Genetics, Institute of Biology, University of Neuchâtel, Neuchâtel, Switzerland; Plant Pathology, Institute of Integrative Biology, ETH Zurich, Zurich, Switzerland

**Keywords:** Amplicon sequencing, long read sequencing, PacBio, Sequel II, multiplexing, pangenome, microbiome, wheat, phyllosphere, *Pseudomonas*, *Zymoseptoria tritici*

## Abstract

High-throughput sequencing technologies have greatly advanced our understanding of microbiomes, but resolving microbial communities at species and strain levels remains challenging. Here, we developed and validated a pipeline for designing, multiplexing, and sequencing highly polymorphic taxon-specific long-read amplicons. We focused on the wheat microbiome as a proof-of-principle and demonstrate unprecedented resolution for the wheat-associated *Pseudomonas* microbiome and the ubiquitous fungal pathogen *Zymoseptoria tritici*. We achieved an order of magnitude higher phylogenetic resolution compared to existing ribosomal amplicons. The designed amplicons accurately capture species and strain diversity outperforming full-length 16S and ITS amplicons. Furthermore, we tracked microbial communities in the wheat phyllosphere across time and space to establish fine-grained species and strain-specific dynamics. To expand the utility of our approach, we generated pangenome-informed amplicon templates for additional key bacterial and fungal genera. Pangenome-informed microbiome profiling enables the tracking of microbial community dynamics in complex environments and overcomes limitations in phylogenetic resolution.

## Main text

High-throughput sequencing technologies (HTS) have revolutionized our understanding of microbiomes revealing insights into structures, functions and dynamics. Microbial community dynamics are influenced by various factors such as the identity of associated hosts, health status, and environmental variables ^1–6^. Targeted amplicon sequencing is the most commonly employed HTS method for microbiome analyses interrogating conserved loci such as 16S ribosomal DNA for prokaryotes and internal transcribed spacers (ITS) for eukaryotes. Amplicon sequencing is widely applicable and overcomes challenges such as low input biomass or contamination by host DNA, making amplicon sequencing particularly suitable for plant microbiome analyses where contaminating DNA is often present in substantial amounts ^7^. In contrast, untargeted metagenomic sequencing and assembly can only provide rich information about microbial communities if the targeted organisms are abundant^7^. A major limitation of amplicon sequencing has been the lack of resolution below the genus level in most studies based on either 16S or ITS ^7,8^.

Long-read sequencing based on PacBio or Oxford Nanopore opened the possibility to sequence long amplicons ^9^. PacBio circular consensus sequencing (CCS) generates highly accurate (>99.9%) reads comparable to Illumina and Sanger sequencing but reaching up to 20 kb in length ^10^. Studies based on PacBio sequencing of full-length 16S or ITS sequences achieved high taxonomic resolution ^8,11–13^. Yet, increased resolution would often be necessary to define functional roles of microbiome components. For example, closely related species of the ubiquitous *Pseudomonas* bacteria can have drastically different effects on plant microbiome functions. Certain *P. syringae* strains are pathogenic, but *P. fluorescens*, *P. protegens*, *P. chlororaphis* or *P. putida* species generally promote plant growth ^14–20^. Closely related *Pseudomonas* strains can also differ significantly in volatiles and metabolites production underpinning distinct niche exploitation in the environment ^19,21–23^. Despite the progress, achieving species and strain-level resolution remains challenging though. Thus, sequencing approaches capturing long and highly polymorphic regions beyond 16S and ITS loci are needed to reveal microbiome diversity at an ecologically relevant resolution. Furthermore, multiplexing large numbers of samples and multiple amplicons is required to tackle complex experimental designs and deep analyses of microbiomes across environments.

Here we present an approach for designing, multiplexing, and sequencing highly polymorphic taxon-specific amplicons using PacBio CCS. Our pipeline allows to track complex bacterial and fungal communities at the species and strain level from environmental samples. As a proof-of-principle, we focus on the microbiome of wheat ^24^. The fungal microbiome of wheat is often dominated by the pathogenic fungus *Zymoseptoria tritici* causing septoria tritici blotch ^25–27^. The bacterial microbiome is dominated by Gammaproteobacteria including the genus *Pseudomonas* both above and below ground ^28,29^. The fungal pathogen *Z. tritici* significantly impacts the composition of microbial communities associated with wheat (Kerdraon et al., 2019). For instance, *Z. tritici* suppresses the host immune system to facilitate colonization by strains of the *P. syringae* group ^30^, while specific *P. fluorescens* strains inhibit the growth of *Z. tritici* ^31^.

In this study, we introduce and validate a pipeline for a novel suite of highly multiplexed amplicons that accomplish an order of magnitude higher phylogenetic resolution compared to existing ribosomal amplicons. As a proof-of-principle, we target the wheat-associated *Pseudomonas* microbiome and the major fungal pathogen *Z. tritici*. We achieve unprecedented species and strain-level resolution for both groups in mixed samples, and we highlight the substantial gains in phylogenetic resolution by tracking strains across the wheat canopy and over time. Furthermore, we show that this pipeline can be applied to other ecologically relevant genera in need of high-resolution amplicons including *Rhizobia*, *Streptomyces*, and *Aspergillus*.

## Results

### Pangenome-Informed design of taxon-specific amplicons

We developed new 3-kb long amplicons to enhance the resolution of bacterial and fungal community profiling while maintaining universal amplification within the targeted group of organisms (Figure 1A). To identify highly polymorphic *Pseudomonas* amplicons, we constructed a comprehensive pangenome of 18 high-quality genomes representing all subgroups of the genus. From this analysis, we identified 28 core regions conserved in the *Pseudomonas* pangenome. These core regions served as candidate regions for primer development. For each core fragment, we designed all possible amplicons ranging from 2.7-3.2 kb resulting in 224 amplicon candidates. Nucleotide diversity in aligned core regions was used as a metric to prioritize amplicon candidates.

**Figure 1:**
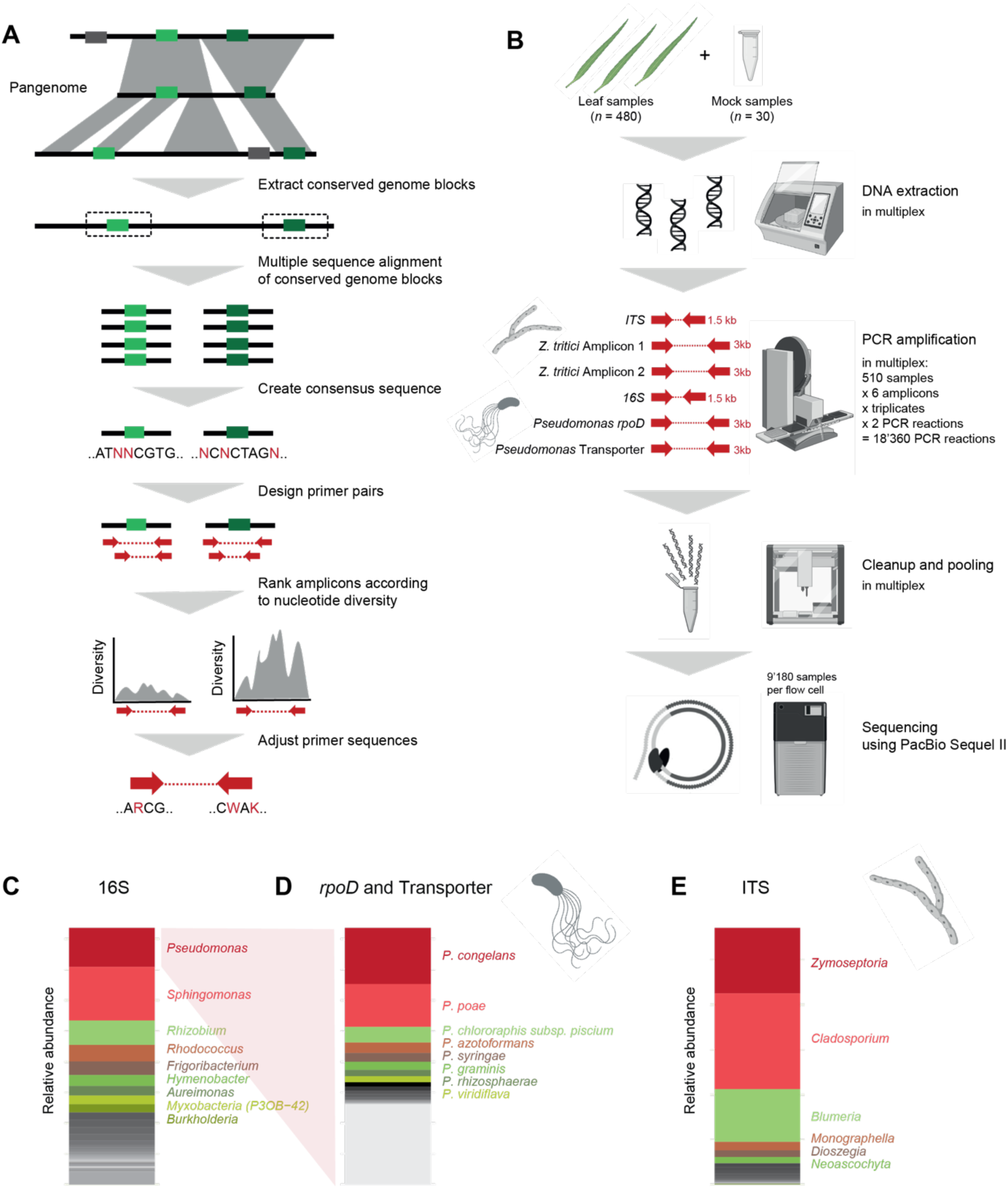
Pangenome-informed design of taxon-specific amplicons. (A) Schematic overview of amplicon design pipeline. Conserved blocks in the *Pseudomonas* and *Z. tritici* pangenomes were extracted. Consensus sequences of the core blocks were used to design amplicons. Amplicon candidates were ranked according to nucleotide diversity, and primer sequences were adjusted. Polymorphic amplicons were tested on reference strains and environmental samples. (B) Overview of multiplexed amplicon evaluation on environmental samples. DNA from mock samples and wheat leave samples were automatically extracted. Two *Pseudomonas*-specific amplicons, two *Z. tritici*-specific amplicons, as well as the full-length 16S and ITS were amplified in each sample and prepared using robotic liquid handling for accuracy and throughput. Reactions were performed in triplicates and barcoded individually using asymmetric M13 barcodes resulting in a total of 18’360 PCR reactions. All reactions were pooled and sequenced in two PacBio Sequel II 8M cells. (C) Relative abundances of bacterial genera on wheat leaves determined by the full-length 16S amplicon. (D) Relative and mean abundances of *Pseudomonas* species on wheat leaves assessed by the *Pseudomonas*-specific *rpoD* and transporter amplicons. (E) Relative abundances of fungal genera on wheat leaves determined by the full-length ITS amplicon.

We selected the ten most polymorphic amplicon candidates for PCR evaluation using both *Pseudomonas* reference strains and naturally colonized wheat leaf samples. Candidate primer sequences were adjusted based on alignment against all available sequences of *Pseudomonas* strains to maximize recovery of the *Pseudomonas* diversity. We allowed primer candidates to include up to five degenerate bases. To reduce the number of sequence variants to be considered in primer candidates, we reduced degenerate positions to match known combinations in *Pseudomonas* genomes. Finally, we selected two amplicons that consistently showed the strongest PCR amplification for further analyses. The first amplicon spanned a locus including the *rpoD* gene, previously utilized for taxonomic classification ^32,33^. The second amplicon encompassed genes encoding an ABC transporter (APE98195.1) and an ABC transporter permease (APE98194.1), respectively. We pursued a parallel approach to identify primers suitable for amplifying the intra-specific diversity of the fungal pathogen *Z. tritici*. We based our analysis on a recently established reference-quality global pangenome for the species ^34^. The two best-performing amplicons were located on chromosomes 9 and 13, respectively.

To evaluate the performance of the two *Pseudomonas*-specific and two *Z. tritici*-specific amplicons, we conducted tests using an extensive mock community of well-characterized laboratory strains. In addition to the new amplicons, we amplified the full-length 16S and ITS loci to contrast performance on the same sample pools. Additionally, we tested the *Pseudomonas* and *Z. tritici*-specific amplicons on a diverse set of wheat leaf samples collected at five timepoints during the growing season at an experiment field site near Zurich, Switzerland (Figure 1B). We optimized DNA extraction methods to maximize bacterial and fungal yield from wheat leaves. Subsequently, we amplified the two selected *Pseudomonas*-specific amplicons and the two *Z. tritici*-specific amplicons in 30 mock and 480 wheat leaf samples. Each reaction was performed in triplicate, resulting in 9,180 amplifications. We tagged each sample with a unique asymmetric M13 barcode combination. Liquid handling was performed with a robot and samples were processed in a fully randomized layout. We sequenced all 9,180 reactions on two PacBio Sequel II flow cells resulting in a total of 1.5M high-quality reads used in all subsequent analyses (Supplementary Table 1).

Our analysis of 480 wheat leaf samples revealed a diverse assembly of bacteria. Based on the full-length 16S amplicon, we found *Pseudomonas* and *Sphingomonas* to be the dominant genera (Figure 1C and Supplementary Figure S1). The *Pseudomonas*-specific *rpoD* and transporter amplicons revealed a diverse assembly of species, primarily belonging to the *P. fluorescens* and *P. syringae* groups (Figure 1D and Supplementary Figure S1). The mycobiome, analyzed using the full-length ITS, was predominantly composed of the fungal genera *Zymoseptoria* and *Cladosporium*. This is broadly consistent with recent findings in wheat fields ^25,27^. However, our dataset showed a higher abundance of *Cladosporium*. This could be attributed to the humid conditions during our sampling season or the limited resolution of the order *Capnodiales* in other amplicon studies on wheat leaves. These studies all analyzed the short ITS1 amplicon (approximately 350bp), compared to our use of the full-length ITS amplicon (approximately 1500bp) (Figure 1E and Supplementary Figure S1).

### Detection limits of taxon-specific amplicons

To assess the performance and detection limits of the new amplicons, we analyzed both defined mock communities and dilution series. For the *Pseudomonas* amplicons, we examined a panel of ten isolates representing the phylogenetic diversity of the genus (Supplementary Table 2). Our results showed that both the full-length 16S and the *Pseudomonas*-specific *rpoD* and transporter amplicons correctly distinguish all isolates (Figure 2A). All isolates consistently amplified for all three amplicons, except for the *P. putida* Leaf58 isolate failing for the transporter amplicon. No genome sequence is available for verification, but amplification failure may be caused by primer mismatches. The *rpoD* and transporter amplicons exhibited clustering of the eight *P. fluorescens* group isolates according to their subclade described by ^35^. While the *rpoD* and transporter amplicons predominantly produced a single amplicon sequence variant (ASV) per culture, the 16S amplicon exhibited multiple ASVs per culture. This can be attributed to the multi-copy nature of the 16S gene ^36^.

**Figure 2:**
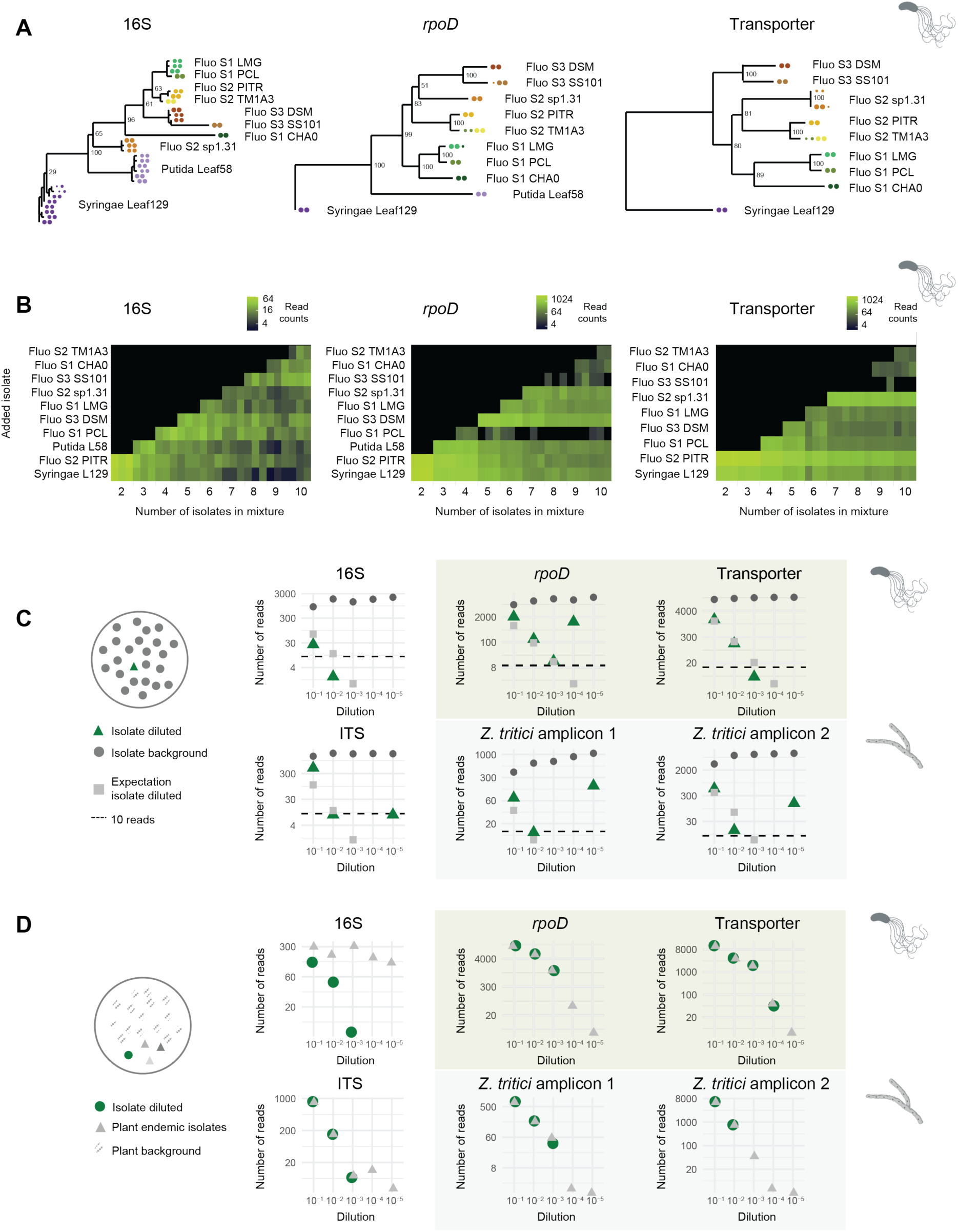
Detection limits of taxon-specific amplicons using mock communities. (A) Phylogenetic trees of amplified sequence variants (ASVs) from ten individual *Pseudomonas* cultures based on the full-length 16S amplicon, the *Pseudomonas*-specific *rpoD* and transporter amplicon. The *Pseudomonas* cultures originate from the following strains: *P. syringae* Leaf129 (Syringae L129), *P. thivervalensis* PITR2 (Fluo S2 PITR), *P. putida* Leaf58 (Putida L58), *P. chlororaphis ssP. piscium* PCL1391 (Fluo S1 PCL), *P. fluorescens* type strain DSM50090 (Fluo S3 DSM), *P. chlororaphis ssP. chlororaphis* type strain LMG5004 (Fluo S1 LMG), *Pseudomonas sP.* 1.31 (Fluo S2 sp1.31), *Pseudomonas sP.* SS101 (Fluo S3 SS101), *P. protegens* CHA0 (Fluo S1 CHA0) and *P. brassicacearum* TM1A3 (Fluo S2 TM1A3). For the isolates from the *P. fluorescens* group, the subclades Fluo S1, S2 and S3 are indicated as defined by Flury et al., 2016. Colors indicate different cultures, and the dot size corresponds to the abundance of the ASVs for each of the two replicates. The transporter amplicon failed to amplify the *P. putida* Leaf58 isolate. (B) Mixed equimolar combinations of the ten *Pseudomonas* cultures showing read counts of the main ASV for each mixture. The three replicates are shown individually. (C) Top panel: Serial dilution series of *P. syringae* Leaf129 (green) mixed into *P. thivervalensis* PITR2 (grey). Read numbers are shown for the full-length 16S, and the *Pseudomonas*-specific *rpoD* and transporter amplicons. The expected read numbers of the diluted *P. syringae* Leaf129 isolate are indicated in light grey and calculated from the background read numbers of *P. thivervalensis* PITR2. Bottom panel: Serial dilution series of *Z. tritici* ST99CH 1E4 (green) in *Z. tritici* ST01IR 48b (grey) isolates. Read numbers are shown for the full-length ITS, and the *Z. tritici*-specific amplicons on chromosomes 9 and 13. The expected read numbers of the diluted *Z. tritici* ST99CH 1E4 isolate are indicated in light grey and calculated from the background read numbers of the *Z. tritici* ST01IR 48b isolate. (D) Top panel: Read numbers of a serial dilution series of *P. thivervalensis* PITR2 (green) mixed into a wheat leaf DNA extract background for the full-length 16S amplicon and the *Pseudomonas*-specific *rpoD* and transporter amplicons. In grey, the total read count of the wheat-endemic bacterial isolates is indicated for the 16S amplicon, respectively the total read count of the wheat-endemic *Pseudomonas* for the *Pseudomonas*-specific *rpoD* and transporter amplicons. Bottom panel: Read numbers of a serial dilution series of *Z. tritici* ST99CH 1E4 (green) mixed into a wheat leaf DNA extract background for the full-length ITS and the *Z. tritici*-specific amplicons. In grey, the total read count of the wheat-endemic fungal isolates is indicated for the ITS amplicon, respectively the total read count of the wheat-endemic *Z. tritici* for the *Z. tritici*-specific amplicons. For all serial dilutions of (C) and (D), dilutions are performed with an initial DNA input of 7.5 ng per isolate.

To evaluate amplicon performance in more complex strain mixtures, we created equimolar mixtures containing two to ten *Pseudomonas* isolates. The 16S, the *rpoD* and the transporter amplicons successfully recovered ASVs from individual isolates even in complex mixtures (Figure 2B). However, the *rpoD* amplicon showed reduced amplification efficiency for the isolate *P. chlororaphis ssP. piscium* PCL1391 and *P. fluorescens* SS101 in the mixture. The transporter amplicon exhibited lower amplification efficiency for the isolate *P. fluorescens* SS101. Furthermore, we observed greater variation in abundance between replicates for the 16S amplicon compared to the *rpoD* and transporter amplicons.

We aimed to achieve very high levels of multiplexing and, as a consequence, recovered relatively low read numbers per sample. To assess detection limits in our experimental setup, we analyzed a two-strain dilution series consisting of the *P. syringae* Leaf129 and *P. thivervalensis* PITR2 isolates (Figure 2C, Supplementary Figure S2, Supplementary Table 3). Both *Pseudomonas*-specific amplicons were able to identify the diluted isolates at a concentration as low as 10^-3^, which corresponds to 7.5*10^-3^ ng input DNA or approximately 1050 cells. On the other hand, the 16S amplicon could detect the diluted isolates only at a slightly higher concentration of 10^-2^, representing 7.5*10^-2^ ng input DNA or approximately 10’520 cells (Figure 2C). Similarly, we performed a comparable analysis for the fungal pathogen *Z. tritici* using a two-strain dilution series comprising *Z. tritici* isolates ST99CH 1E4 and ST01IR 48b (Figure 2C, Supplementary Figure S2, Supplementary Table 3). The ITS and *Z. tritici*-specific amplicons successfully detected diluted strains down to a concentration of 10^-2,^ corresponding to 7.5*10^-2^ ng input DNA or approximately 1720 cells. Polyphenols contained in plant DNA extracts can inhibit amplification. To test for this, we replicated the dilution series by diluting reference isolates with leaf samples. We obtained similar amplification yields from culture samples compared to plant samples indicating that the remaining plant extracts did not significantly affect the amplification (Figure 2D, Supplementary Figure S2, Supplementary Table 3).

### Discriminant power of amplicons to resolve strain genotypes

To assess gains in phylogenetic resolution of the wheat microbiome, we compared the *Pseudomonas* and *Z. tritici* amplicons to 16S and ITS amplicons, respectively. Analyzing 480 wheat leaf samples, Pseudomonas-specific amplicons revealed in total 933 and 538 ASVs at the *rpoD* and transporter locus, respectively. In contrast, the 16S amplicon revealed only 86 ASVs matching the genus *Pseudomonas* (Figure 3A and B, Supplementary Figure S3). This represents a three-fold (2.7X for *rpoD* and 3.3X for the transporter) increase in ASVs for the *Pseudomonas*-specific amplicons compared to the 16S and based on relative number of reads per amplicon (Figure 3B).

**Figure 3:**
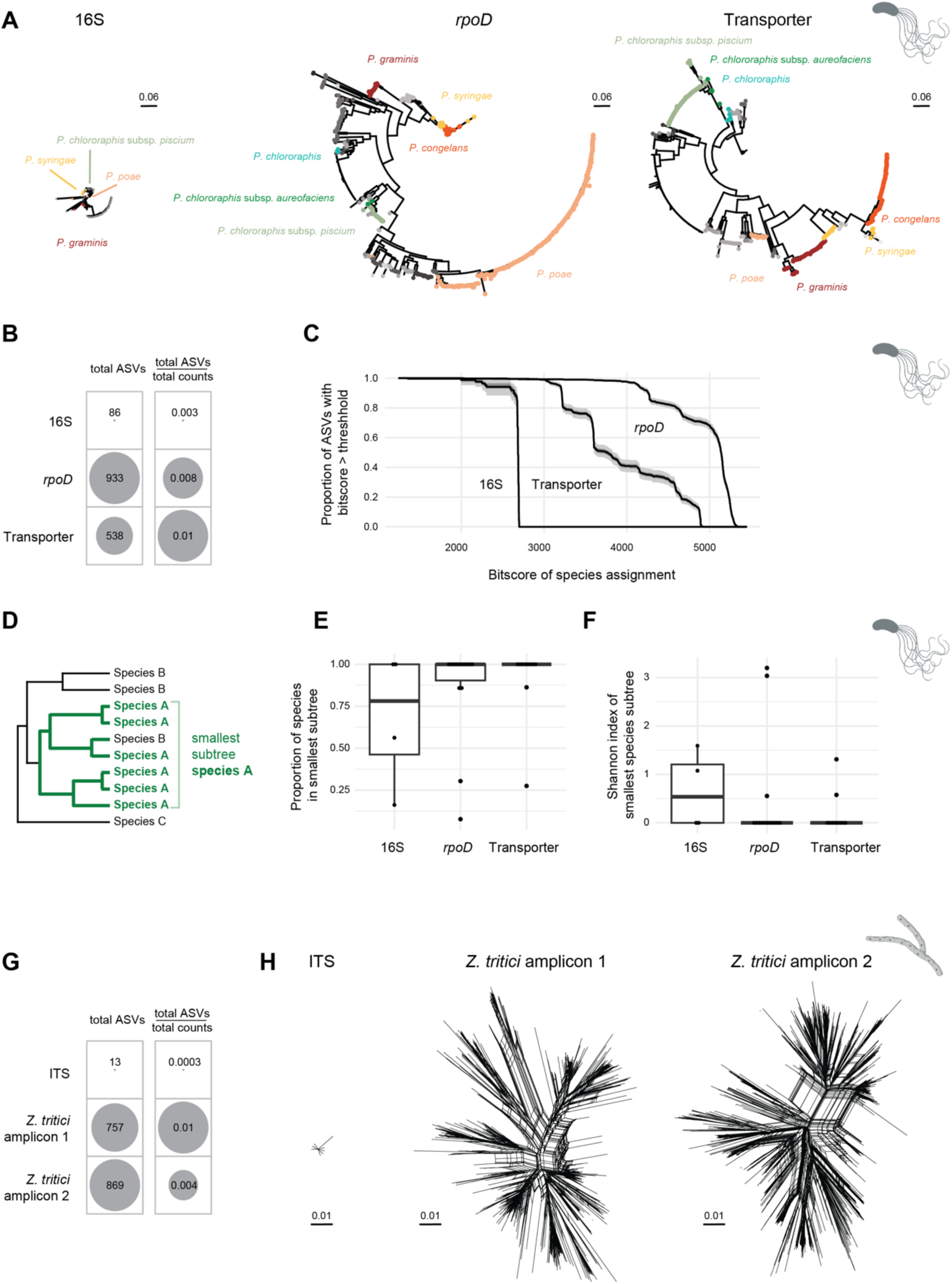
Phylogenetic resolution of taxon-specific and ribosomal DNA amplicons. Taxon-specific amplicon analysis based on 480 wheat leaf samples. (A) Phylogenetic trees of *Pseudomonas* amplified sequence variants (ASVs) of the full-length 16S amplicon as well as the *Pseudomonas*-specific *rpoD* and transporter amplicons. ASVs are colored by *Pseudomonas* species. (B) Total number of ASVs detected by the full-length 16S amplicon, and the *Pseudomonas*-specific *rpoD* and transporter amplicons relative to the total read count per amplicon. (C) Proportion of ASVs with a BLASTn bitscore above a threshold for *Pseudomonas* species assignment. The 50% and 95% confidence intervals based on permutations are shown in grey. (D) Identification of the smallest subtree containing >90% of the ASVs for each *Pseudomonas* species shown in (E) and (F). Only species with >5 ASVs were considered. (E) Proportion of species in the smallest subtree containing >90% of the species ASVs assigned to specific species. (F) Shannon diversity indices of the subtree species composition for the smallest subtree containing >90% of the ASVs assigned to specific species. (G) Total number of ASVs detected by the full-length ITS amplicon, and *Z. tritici*-specific amplicons on chromosome 9 and 13 relative to the total read count per amplicon. (H) Phylogenetic networks (Splitstree) of *Z. tritici* ASVs identified using the full-length ITS amplicon, and the *Z. tritici*-specific amplicons.

We assigned all ASVs to *Pseudomonas* species using 1071 available genomes from nine different groups for BLASTn analyses (Supplementary Table 4). The *Pseudomonas*-specific amplicons produced significantly better matches for species assignment compared to 16S sequences (Figure 3C). To assess the consistency of species assignments, we extracted the smallest subtree containing >90% of the ASVs for each *Pseudomonas* species and examined the proportion of the species within this subtree, as well as the Shannon diversity index of the ASVs matching each species (Figure 3D-F). *Pseudomonas*-specific amplicons showed higher proportions and lower Shannon diversity indices compared to the 16S as expected for more accurate species assignments. For this analysis we considered four species for the 16S, 14 for *rpoD*, and ten for the transporter amplicon, where the low species number for 16S stems from the requirement of at least five detected ASVs per species (Supplementary Figure S4). Among the species resolved by the *rpoD* and the transporter amplicons, but failing with the 16S, were subspecies of *P. chlororaphis* and various species from the *P. syringae* group (Figure 3A and Supplementary Figure S3).

For *Z. tritici*, we found 869 and 757 ASVs for the chromosome 9 and 13 amplicon, respectively, compared to 13 ASVs obtained by ITS (Figure 3G and H). Based on the total number of reads per amplicon, this represents a more than tenfold increase of ASVs for the *Z. tritici*-specific amplicons. These findings indicate a remarkably high strain diversity within a single field in agreement with previous studies ^37,38^.

### High-resolution tracking of natural plant colonization by pseudomonads

A major aim of high-resolution microbiome analyses is to resolve strain-level interactions. To assess the power of the newly developed amplicons, we performed a hierarchical sampling of 480 samples across space and time tracking expected microbiome shifts in a wheat field. Specifically, sampling was conducted at five different timepoints during the wheat growing season, from May (first node appearance) to July (prior to harvest). At each timepoint, leaves were sampled at three different canopy heights to capture developmental patterns of the plant.

We first examined whether *Pseudomonas* species abundance patterns were consistently reproduced by both the *rpoD* and transporter amplicons. For this, we calculated correlation coefficients of *Pseudomonas* species abundance across samples and between the two amplicons (Figure 4A and Supplementary Figure S5). We compared correlation values to null expectations based on permutations. We observed that 68% and 77% using Spearman and Pearson correlations, respectively, of the species correlations were higher than the 95% confidence interval of the null expectation.

**Figure 4:**
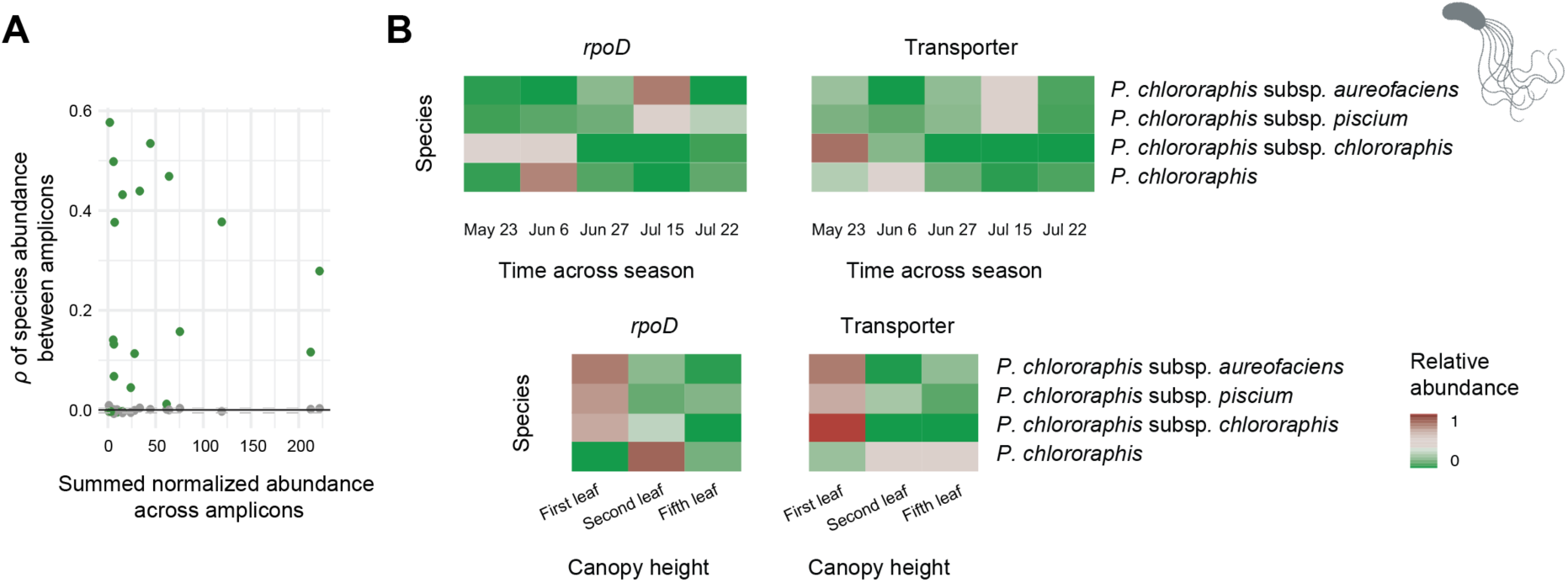
Differential abundance of wheat phyllosphere *Pseudomonas* species across season and canopy height. (A) Spearman correlation of *Pseudomonas* species abundance across samples between the *Pseudomonas*-specific *rpoD* and transporter amplicons. Correlation values were compared to null expectations based on permutations with the grey dotted line indicating the 95% confidence interval of the null expectation. (B) Relative abundances of *P. chlororaphis* subspecies across time and canopy heights revealed by the *Pseudomonas*-specific *rpoD* and transporter amplicons.

Next, we investigated species abundance changes across time and space. We identified different subspecies of *P. chlororaphis* exhibited variable abundance patterns throughout the season and across canopy heights in both amplicons (Figure 4B). *P. chlororaphis* subsP. *aurofaciens* and subsP. *piscium* were most abundant in July, whereas *P. chlororaphis* and *P. chlororaphis* subsP. *chlororaphis* were most abundant in May and June. *P. chlororaphis* subsP. *aurofaciens*, subsP. *piscium* and subsP. *chlororaphis* were most prominent at the bottom of the canopy, whereas *P. chlororaphis* was most abundant on upper leaves.

### Within-Species diversity of a major wheat pathogen

Genetic diversity within pathogen species can underpin rapid breakdowns of fungicide efficacy or host resistance ^39,40^. How genotypic diversity changes over pathogen life cycles remains largely unknown though ^41,42^. Here, we examined changes in genotypic diversity across the epidemic phase for the fungal pathogen *Z. tritici* based on species-specific amplicons. The two *Z. tritici*-specific amplicons showed consistent numbers of ASVs per leaf throughout the season, indicating minimal turnover in pathogen diversity within a single epidemic phase (Figure 5A). Similarly, the number of ASVs did not differ significantly across different canopy heights (Figure 5A). Tracking individual ASVs across the season, we observed two distinct groups based on their abundance patterns (Figure 5B). The first group consisted of a small number of highly abundant strains persisting throughout the entire growing season. In contrast, the second group consisted of numerous strains that are predominantly scarce and only detected at specific timepoints.

**Figure 5:**
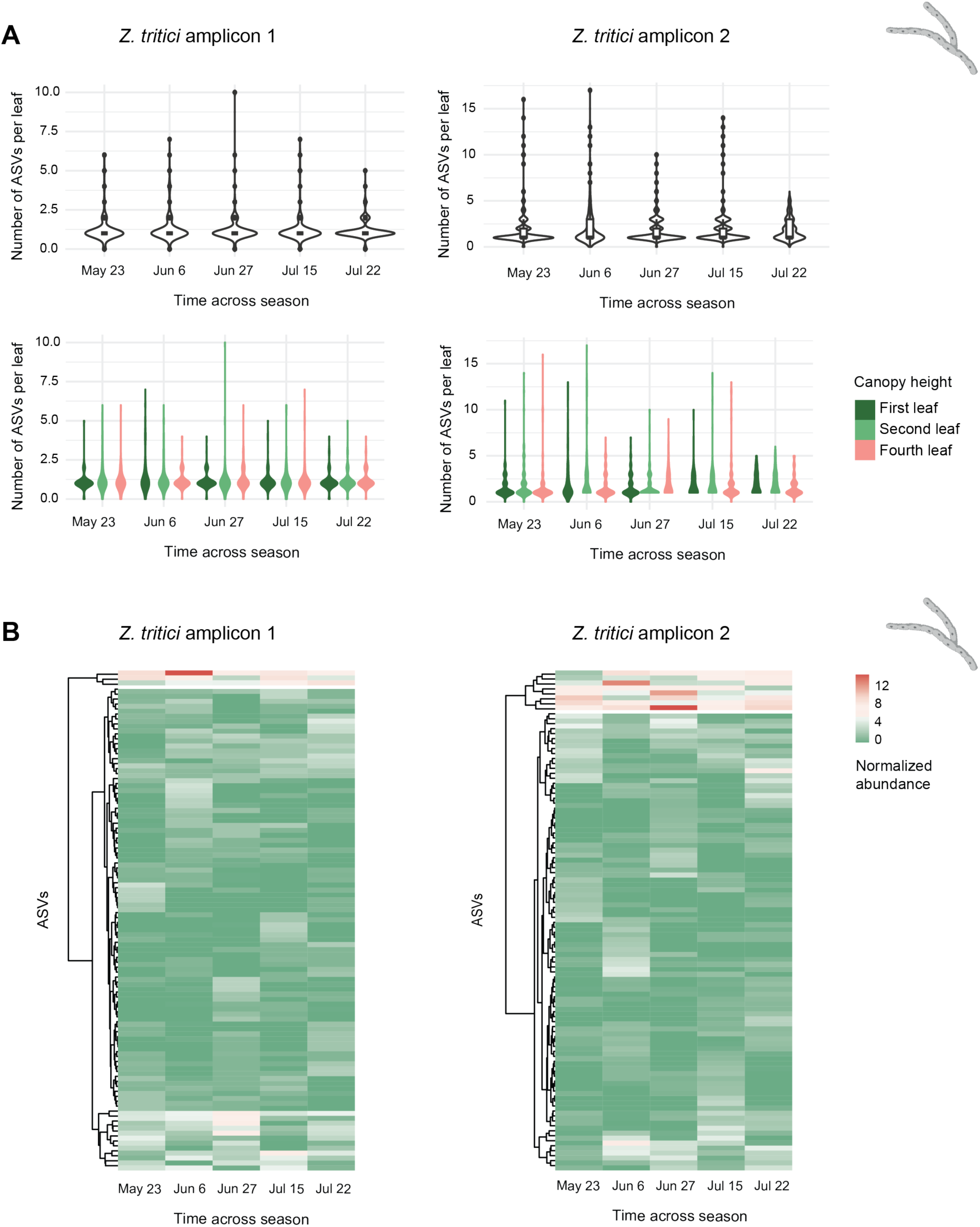
Genotypic diversity of the wheat phyllosphere pathogen *Z. tritici* across season. (A) Number of unique *Z. tritici* amplicon sequence variants (ASVs) per leaf across timepoints based on the *Z. tritici*-specific amplicons on chromosome 9 and 13. The same analysis is shown highlighting canopy differentiation. (B) Heatmap of the 100 most abundant *Z. tritici* ASVs clustered based on their abundance pattern across the season.

### Pangenome-Informed Amplicon Templates for Additional Taxa

We assessed the potential to generate high-resolution amplicons for other ecologically relevant taxa. We used the same principle as for *Pseudomonas* to define a representative set of high-quality genomes to construct a pangenome to search for core regions conserved within the taxon. Such regions can then be systematically examined for potential amplicon sequences. We generated pangenomes for *Rhizobia* and *Streptomyces* following a systematic procedure to allow for comparison (Figure 6 and Supplementary Figure S6). We also investigated the important environmental fungal species *Aspergillus fumigatus* causing opportunistic human infections. The pangenome analyses defined 22 core regions for *Rhizobia*, 515 for *Streptomyces*, and 5,107 for *A. fumigatus*, respectively. These findings indicate genomic regions that are suitable for primer design (Figure 6 and Supplementary Figure S6). The relatively small number of candidate core regions for *Rhizobia* correlates with its greater 16S rRNA gene diversity compared with *Streptomyces* and *Pseudomonas*. Additionally, all pangenome genes were categorized into orthogroups based on protein homology. Our analysis reveals that the proportion of orthogroups present in all isolates of the pangenome varies from 16% to 84% among the analyzed taxon-specific pangenomes. This range reflects the extent of diversity within the pangenome that our method can accommodate (Figure 6 and Supplementary Figure S6).

**Figure 6:**
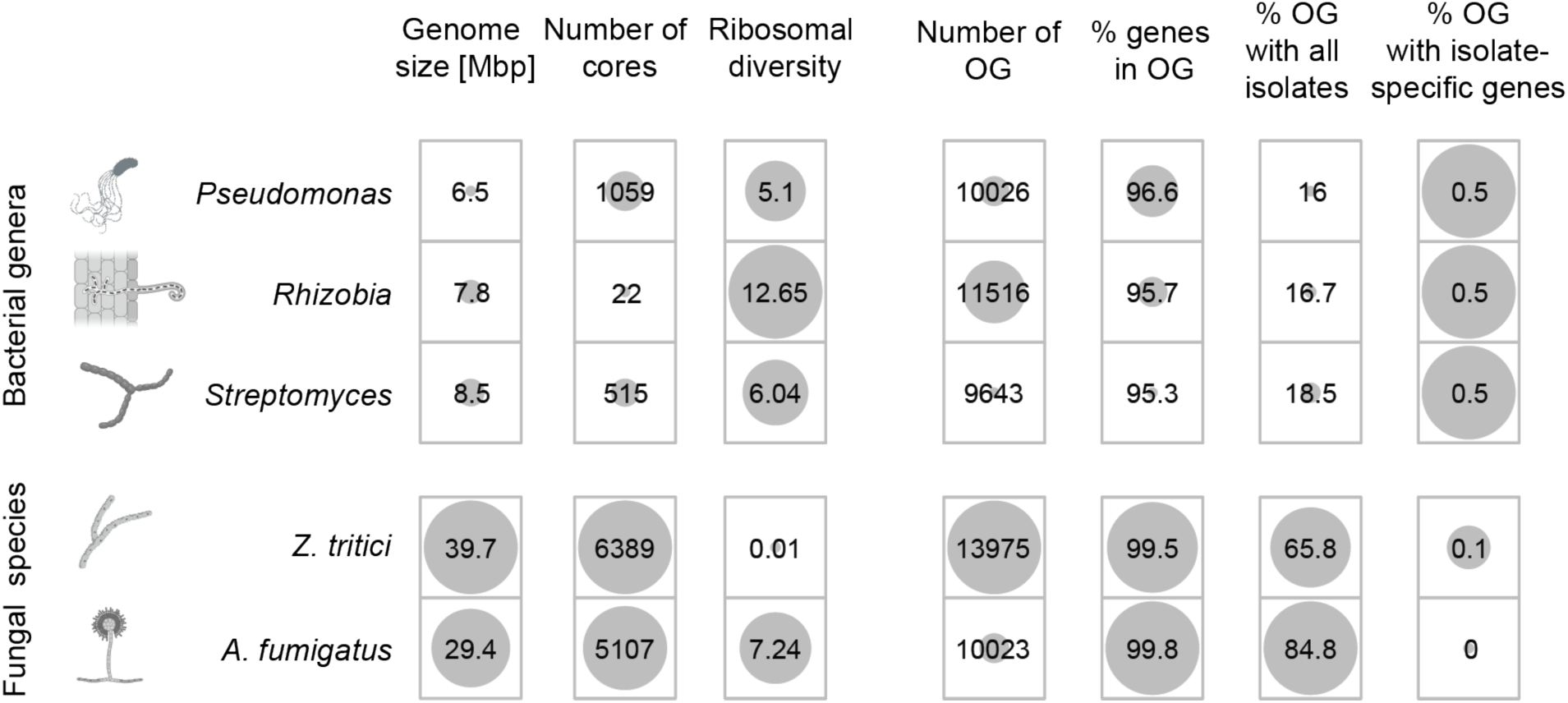
Pangenome and amplicon templates for the bacterial genera *Pseudomonas*, *Rhizobia*, and *Streptomyces*, as well as the fungal species *Z. tritici* and *A. fumigatus*. Pangenomes were each generated based on 19 representative genomes. For each pangenome, the average genome size in Mbp, the number of candidate core segments and the length-normalized ribosomal nucleotide diversity of the 16S or ITS regions is indicated. Genes were grouped into orthogroups (OG) based on protein homology for each pangenome. The total number of orthogroups, the percentage of genes included in orthogroups, the percentage of orthogroups that contain all isolates, and the percentage of orthogroups that contain isolate-specific genes are shown for each species group.

## Discussion

We demonstrate that limitations in phylogenetic resolution of microbial community profiling can be overcome using systematically designed taxon-specific 3-kb amplicons. We find that the new loci provide species and strain-level insights into subsets of bacterial and fungal communities coexisting in the plant microbiome exceeding full-length 16S or ITS amplicons by up to an order of magnitude.

The pangenome-informed design of amplicons for pseudomonads was optimized to capture diversity specifically and exclusively in this ubiquitous group of bacteria. Compared to full-length 16S amplicons, the new amplicons provide consistently higher confidence in species assignments across clades. The confidence in species assignments stems from the high degree of species resolved to monophyletic clades for most *Pseudomonas* species. This enhanced resolution enabled us to differentiate *Pseudomonas* at the subspecies level, which is not feasible using 16S alone. We also focused on the dominant wheat phyllosphere pathogen *Z. tritici* and achieved a strain-level resolution inaccessible with existing single-locus genotyping approaches. The recovered diversity reflects genotyping resolution typically only achievable by whole-genome sequencing ^37,38^.

We evaluated the discriminant power of the new bacterial and fungal amplicons by tracking microbial communities in the wheat phyllosphere. We contrasted the resolution to the universal barcoding loci 16S and ITS in samples covering different time points and canopy heights. The increased resolution indeed revealed species and strain-specific changes across time and canopy height inaccessible by ribosomal barcoding datasets. One noteworthy finding was the differential abundance patterns observed for the *P. chlororaphis* subspecies. Subspecies of *P. chlororaphis* have garnered agricultural interest due to their biocontrol potential, with several strains being utilized in commercial formulations ^19,43^. This potential is attributed to the production of antimicrobial metabolites, notably phenazines, which are not only linked to microbial antagonism but also plant defense upregulation ^19,43^. However, phenazine expression and regulation vary among *P. chlororaphis* subspecies. Therefore, monitoring *P. chlororaphis* at the subspecies level is crucial for analyzing antagonistic behavior in the field and prioritizing the most promising biocontrol candidates. The assay developed here is the first to facilitate this. Additionally, we were able to resolve genotypic diversity in a highly diverse fungal pathogen and track changes throughout a single epidemic season. The virulence of different *Z. tritici* isolates varies greatly, making it interesting to track the genotypic diversity across the epidemic season^44,45^. Despite the overall large genotypic diversity (>700 ASVs within a single field), we identified a small set of genotypes that dominated the entire season, which can be informative for control strategies. Our data would also allow us to infer strain-specific bacterial-fungal interactions based on abundance data, to prioritize biocontrol candidates antagonizing specific pathogen isolates.

The replicated monitoring of *Pseudomonas* and plant pathogen diversity with two independent amplicons each enabled us to assess the reproducibility of species and strain identification as well as abundance estimates. Certain *Pseudomonas* species were over- or underrepresented in a particular amplicon and across samples. Such discrepancies could stem from differences in primer binding efficiency and variations in amplification success due to amplicon sequence variability ^46^. Our findings align with previous studies investigating reproducibility of different 16S amplicons ^47^. This underscores the importance of implementing controls in quantitative microbiome studies. The availability of two independent 3-kb amplicons for *Pseudomonas* in addition to a control by the 16S provides a powerful toolset to track the reproducibility of microbiome community assessments across samples and taxonomic groups.

In our proof-of-principle demonstration, we multiplexed nearly 10,000 reactions in a single sequencing run. This achievement was made possible by recent improvements in Sequel II. The quality of the sequences was high enough to recover sequences at an impressive rate using asymmetric barcoding, which allowed us to multiplex nearly 10,000 samples with the standard set of barcodes. Therefore, the level of multiplexing in this workflow can be determined solely by the number of reads desired per sample. With taxon-specific amplicons, the sample complexity is lower, making higher multiplexing desirable. Additionally, we enhanced the efficiency and reproducibility of sample handling through robotic liquid handling.

The high quality of PacBio HiFi reads enables detailed microbial community profiling using long amplicons. With many high-quality genomes available, we can leverage pangenomes for more integrative analyses of genetic diversity within taxonomic groups and identify the most informative amplicons. Despite the increasing use of pangenomes to analyze genetic diversity, they have not been utilized for amplicon design to the best of our knowledge ^48–50^. With the rapidly growing number of genomes, this approach has broad applicability. We demonstrated this by calculating the pangenome for the bacterial groups *Rhizobia* and *Streptomyces*, as well as for the fungal species *A. fumigatus*, and identified an array of candidate regions suitable for primer design. A recent study on *Streptomyces* taxonomy found that 16S rRNA sequence variation does not reliably delineate *Streptomyces* species ^51^. The study advocates for alternative markers and suggests a reclassification of *Streptomyces* taxonomy, historically based on 16S data. *Rhizobia* is a highly diverse bacterial group characterized by its ability to fix atmospheric nitrogen. Species delineation often involves multilocus sequence analysis using housekeeping genes ^52–54^. Longer, high-resolution amplicons could track community dynamics in soil and root nodules, revealing species competitiveness in nodule occupancy. *A. fumigatus* is both an opportunistic human fungal pathogen as well as an environmental saprobe. Despite extensive genomic analysis of the species, the diversity in environmental versus clinical niches is still poorly understood ^55,56^. Similar to other fungal species, metagenomic analysis for *A. fumigatus* remains challenging due to the small fraction of fungal DNA in samples and the significantly larger genome compared to bacteria ^57^. High-resolution amplicons could provide a suitable approach to gain strain resolution of environmental and clinical samples, enhancing our understanding of strain competitiveness across different environments.

Overall, our approach yielded similar costs per high-quality sequence as the commonly used 16S and ITS Illumina-based amplicon sequencing. This gain in throughput opens cost-effective avenues to explore microbial community dynamics with complex experimental designs and in heterogeneous environments. Furthermore, designing, optimizing, and multiplexing long amplicons for other ecologically relevant taxa is clearly in reach with the availability of high-quality genomes. In conclusion, our work shows that long-read amplicon sequencing based on purpose-designed taxon-specific amplicons overcomes limitations in phylogenetic resolution associated with ribosomal amplicon sequencing.

## Methods

### Wheat leaves sampling

We analyzed eight elite European winter wheat (*Triticum aestivum*) varieties sampled at five different timepoints over the growing season (i.e. cultivars Aubusson, Arobase, Lorenzo, CH Nara, Zinal, Simano, Forel and Titlis on the 23.05.2019, 06.06.2019, 27.06.2019, 15.07.2019 and 22.07.2019, respectively). The wheat developed from the growth stage Feekes 7.0 until 11.4 ^58^. Two biological replicates of the wheat panel were grown in two complete block designs separated by approximately 100m at the field phenotyping platform site of the Eschikon Field Station of the ETH Zurich, Switzerland (coordinates 47.449°N, 8.682°E). The cultivars were grown in plots of 1.2-by-1.7-m, with the genotypes arranged randomly within each block. No fungicides were applied. For each cultivar, block and timepoint, two plants were collected. From each plant three leaves were collected: the bottom leaf touching the ground (first leaf), the lowest leaf not touching the ground (second leaf) and then the flag leaf (fourth leaf). Each leaf was immediately stored in a plastic foil to avoid contamination, stored at 4°C overnight before processing. In total, 480 leaves were collected (8 cultivars x2 blocks x5 timepoints x2 plants x3 leaves).

### Mock community creation

We established three sets of mock communities using ten different *Pseudomonas* strains and two *Z. tritici* isolates (Supplementary Table 2). The first set was composed of a ten-fold serial dilution series of the DNA from two *Pseudomonas* isolates (*P. syringae* Leaf129 and *P. thivervalensis* PITR2) and two *Z. tritici* isolates (ST99CH 1E4 and ST01IR 48b), up to a dilution of 10^-5^. The DNA input per sample was 15 ng, with 7.5 ng DNA each from *Pseudomonas* and *Z. tritci* for the undiluted sample. For the 10^-1^ dilution, we used 0.75ng DNA for the *Pseudomonas* and *Z. tritici* isolate that was to be diluted, together with 6.75 ng of the background *Pseudomonas* and *Z. tritici* isolate respectively. The second set consisted of a ten-fold serial dilution series up to 10^-5^, where *P. thivervalensis* PITR2 and ST99CH 1E4 were diluted in a leaf DNA extraction sample. We used two different leaf DNA extracts, Leaf 1 (Timepoint 2, Cultivar Zinal, Plot2, Plant 2, Leaf height: top) and Leaf 2 (Timepoint 3, Cultivar Titlis, Plot 2, Plant 2, Leaf height: top). The DNA input per sample was 15 ng, starting with 1.5 ng of isolate and 8.5 ng of leaf DNA for the 10^-1^ dilution. For the third set, we combined increasing numbers of *Pseudomonas* isolate DNA extracts (from two to ten isolates) in equimolar concentrations. The total DNA input per mixture was 15 ng, adding 7.5 ng per isolate for the two-strain mixture and 1.5 ng for the 10-strain mixture. To estimate the number of cells per input DNA, we based our calculations on the average genome size of *Pseudomonas* (6.5 Mbp) and *Z. tritici* (39.7 Mbp), respectively. We used a conversion factor of 660 daltons per base pair for these calculations.

### Pangenome construction

For pangenome construction, we used Panseq with the following settings: fragmentationSize = 5000, minimumNovelRegionSize = 500, novelRegionFinderMode = no_duplicates, percentIdentityCutoff = 60, runMode = pan, storeAlleles = 1, allelesToKeep = 2, frameshift = 1, overwrite = 1, maxNumberResultsInMemory = 500, blastWordSize = 11, nucB = 200, nucC = 65, nucD = 0.12, nucG = 90, nucL = 20, cdhit = 0, sha1 = 0 ^59^. To create the *Pseudomonas* pangenome, we selected 19 high-quality genomes representing all subgroups of the genus ^32,33^ (Supplementary Table 5). For the *Z. tritici* pangenome, we used 19 global reference isolates used for the pangenome analysis of Badet et al. 2020 (Supplementary Table 6). For the *A. fumigatus* pangenome creation, we used all publicly available genomes of the pangenome study of Barber et al. 2021 (n=253, Supplementary Table 7). For the *Rhizobia* pangenome we used the genomes of the pangenome analysis of Yang et al. 2020 (n= 84, Supplementary Table 8). For the *Streptomyces* pangenome, we based the genome selection on Kieper et al. 2023. They clustered all available Streptomyces genomes into 186 distinct species based on average nucleotide identity. For our pangenome construction, we aimed to use one genome per species, but filter for complete genomes or chromosome-level assemblies, leaving us with 97 genomes for pangenome construction (Supplementary Table 9). Bacterial plasmid sequences were removed for all pangenome analysis. Furthermore, to be able to better compare the *Rhizobium*, the *Streptomyces* and the *A. fumigatus* pangenome characteristics to the *Pseudomonas* and *Z. tritici* pangenomes, we subset the genome sets to 19 genomes each for a second pangenome construction based on the same number of genomes (Supplementary Table 6-9). Regions that were present in >50% of the pangenome isolates were selected as “core” for further analysis of the amplicon primer design. We used this relaxed criterion to be able to scan divergent regions for amplicons potentially revealing the highest resolution among strains. Supplementary Table 10 details the *Pseudomonas* pangenome core regions, and Supplementary Table 11 details the *Z. tritici* pangenome core regions. Orthologues genes of all pangenomes were identified Orthofinder v2.5.5 with default parameters (Emms and Kelly, 2019).

### Amplicon primer design

For each pangenome core fragment of the *Pseudomonas* and the *Z. tritici* pangenome, we created a multiple sequence alignment using muscle v.3 with default parameters ^61^. From each core multiple sequence alignment, we created a consensus sequence using the EMBOSS v.6 cons function, setting the identity to ten and plurality to 0.8 ^62^. We then ran Primer3 v. 2.5 on each core consensus sequence^63^. We used the following Primer3 settings: PRIMER_PICK_LEFT_PRIMER=1, PRIMER_PICK_INTERNAL_OLIGO=0, PRIMER_PICK_RIGHT_PRIMER=1, PRIMER_OPT_SIZE=20, PRIMER_MIN_SIZE=18, PRIMER_MAX_SIZE=22, PRIMER_PRODUCT_SIZE_RANGE=1500-3100, PRIMER_NUM_RETURN=8, PRIMER_EXPLAIN_FLAG=1. For each primer pair, we cut out the respective amplicon from the multiple sequence alignment using the EMBOSS v. 6 extractalign function ^62^. To evaluate amplicon diversity, we calculated the nucleotide diversity of each amplicon multiple sequence alignment within the pangenome. For this, we first converted the multiple sequence alignment to vcf format using snp-sites ^64^, and then vcftools v0.1.16 with the options –sites-pi –haploid to calculate the nucleotide diversity ^65^. For both *Pseudomonas* and *Z. tritici*, we selected the ten amplicons with the greatest summed nucleotide diversity for further evaluation. Each *Pseudomonas* candidate primer pair was then blasted against all available *Pseudomonas* genomes from the ncbi nucleotide collection using BLASTn to ensure matches in all genomes ^66^. All hits were aligned using MAFFT v7.427 with the option –auto ^67,68^. For *Z. tritici*, we analyzed each candidate amplicon against the worldwide collection of 1109 *Z. tritici* isolates ^69^. Here, amplicon sequences were extracted from variant call files (i.e. vcf) using bedtools filter ^70^. For both *Pseudomonas* and *Z. tritici* amplicon evaluation, we assessed allele frequencies at every position of the multiple sequence alignment using base R functions ^71^. We manually adjusted primer sequences to account for the specific combinations of alleles necessary to amplify all known *Pseudomonas,* or *Z. tritici* sequences, respectively.

All ten *Pseudomonas* and ten *Z. tritici* primer candidates were tested on reference cultures of *Pseudomonas* and *Z. tritici*, respectively, as well as on naturally infected wheat leaf samples. The two *Pseudomonas* and two *Z. tritici* primer pairs with most consistent amplification success were retained for further steps.

### Sample homogenization, DNA extraction, amplification, Pooling and cleanup

Leaves were lyophilized for 48 h and weighed. Then, the complete leaves were homogenized using 0.5 mm and 0.2 mm zirconium beads in the Bead Ruptor bead mill homogenizer (OMNI) using the following settings: speed 5.00, number of cycles 2, time of cycle 1:00, time distance between cycles 1:00. DNA extraction was performed with automated magnetic-particle processing using the KingFisher Flex Purification Systems (Thermo Scientific). To enhance the DNA extraction of fungal and bacterial DNA, lyticase and lysozyme was added to the first lysis step with PVP buffer. Specifically, for 10mg dry leaf mass 3.9 µl lyticase (200,000 U/mg, diluted to 6.5 mg/ml), 3.9 µl lysozyme (22,500 U/mg, diluted to 10 mg/ml) and 98µl PVP lysis buffer was added, and samples were incubated at 55°C for 30min. Then, 3.9µl proteinase K (30 U/mg, diluted to 10mg/ml) for 10mg dry mass was added and incubated at 55°C for 30min. From each sample, 150µl of clear lysate was transferred to an empty binding plate (KingFisher Flex, Thermo Scientific). For each sample, 360 µl PN binding buffer, 30 µl well suspended Sbeadex beads were added. Using the KingFisher Flex, the first washing step was performed using 400 µl PN1 buffer per sample, then a second wash using 390 µl buffer PN1 with 10µl RNase A (diluted to 10mg/µl in water) and a third wash using 400 µl PN2 buffer. Each sample was eluted in 100 µl nuclease-free water. The DNA concentration was measured using the Spark Microplate reader (Tecan). Then, DNA concentrations were diluted to 5 ng/µl using the Liquid Handling Station (BRAND). PCR reactions were pipetted using the Mosquito HV liquid handling robot (SPT Labtech). The first amplicon PCR reaction was performed in a 15 µl reaction volume. Specifically, 7.5 µl KAPA HiFi HotStart ReadyMix (2x), 1.5µl forward primer (3 µM), 1.5ul reverse primer (3 µM), 3 ul DNA (5 ng/µl) and 1.5 µl HPLC water were combined. Primer sequences and cycling protocols are documented in Supplementary Table 13 and 14. All primers were synthesized by IDT (Integrated DNA Technologies, Coralville, IA). *Pseudomonas* and *Z. tritici*-specific amplicon PCR products were diluted 1:5, PCR products from 16S and ITS 1:10. The second barcoding PCR reaction was performed in a 25 µl reaction volume. Specifically, 12.5 µl KAPA HiFi HotStart ReadyMix (2x), 2.5 µl M13 forward barcode (3 µM), 1.5 µl M13 reverse barcode (3 µM), 2 µl diluted PCR product and 5.5 µl HPLC water were combined. Barcode sequences are available in Supplementary Table 15. Samples were pooled by amplicon taking 1.5 µl from each barcoded product. Each amplicon pool was cleaned using AMPure XP beads using a bead ratio of 0.8x.

### Library preparation and sequencing

Library preparation and sequencing was carried out at the Functional Genomics Centre Zurich (FGCZ). Two SMRTbell libraries were prepared for each amplicon length using the SMRTbell prep kit 3.0. One for the long 3-kb amplicons, one for the 1.5kb 16S and ITS amplicons. Size selection was performed using BluePippin (Sage Science) with a 0.75% dye-free cassette for each library. PacBio sequencing was performed on a Sequel II machine with the SMRT 8M cell. Two sequencing runs were performed, the first one using SMRT Link version 10.1 and the second one SMRT Link version 11.1. The first run encompassed all leaf samples, and the second one all mock samples.

### PacBio raw read processing

CCS were extracted from raw reads using the ccs software from the bioconda package pbccs provided by the manufacturer (Pacific Biosciences). For the first run pbccs v. 6.0.0 was used and for the second run v. 6.3.0. We split the CCS reads by barcodes using the software lima 2.0.0 (Pacific Biosciences) with the following parameters lima –log-level INFO –per-read –min-passes 0 –split-bam-named –ccs –different -A 1 -B 3 -D 2 -I 2 -X 0. We assigned the reads to the respective amplicons using BLASTn assignments to reference amplicon from the *P. fluorescens* CHA0 and *Z. tritici* isolate 1E4 ^66^. All six reference amplicons were blasted against each read and reads were then assigned to the reference hit with the lowest e-value, the highest length and the highest identity. We removed primer sequences using cutadapt v. 3.4 with the following parameters: cutadapt - a FORWARD_PRIMER_SEQ…REVERSE_PRIMER_SEQ –discard-untrimmed –revcomp ^72^. Primers were treated as linked, i.e. reads without primers at both ends were discarded. The R package dada2 v.

1.28.0 was used to infer ASVs for each amplicon separately ^73^. In the following, dada2 steps for each amplicon are described. Reads were filtered and trimmed using the function filterAndTrim with the parameters minLen=minLength, maxLen=maxLength, rm.phix=FALSE, maxEE=2, qualityType = "FastqQuality", multithread=TRUE. Length ranges for each amplicon are described in Supplementary Table 9. Reads were dereplicated using the function derepFastq with the parameters verbose=TRUE, qualityType="FastqQuality". Error models were estimated with the function learnErrors with the parameters errorEstimationFunction=dada2:::PacBioErrfun, BAND_SIZE=32, multithread=TRUE. Reads were denoised using the function dada with the parameters BAND_SIZE=32, multithread=TRUE, pool=FALSE. Importantly, we did not used the option pool=TRUE, as this introduced spurious ASVs in the reference cultures of the mock community. ASV sequence tables were generated using the function makeSequenceTable. To remove chimeras from the sequence table, the function removeBimeraDenovo with parameters method="consensus", minFoldParentOverAbundance=3.5, multithread=TRUE, verbose=TRUE was used. Supplementary Figure S7 shows the read tracking through the dada2 pipeline for each amplicon.

### Taxonomic classification

We assigned 16S reads using the dada2-formatted Silva database v. 138 ^74^ and ITS reads to the UNITE database v. 8.3 ^75^. To perform taxonomy assignment, we used the function assignTaxonomy from the dada2 package. As the UNITE database comprises mostly ITS1-ITS2 reference sequences and only few full-length ITS-LSU sequences, we truncated all ITS reads to the ITS1-ITS2 fragment for assignment. We blasted the reads against the ITS1-ITS2 fragment of *Z.tritici* strain S-46 (KT336200.1) and used the hit coordinates to truncate reads with the seqkit software function subseq ^76^. We assigned *Pseudomonas* reads of the *rpoD*, transporter and 16S amplicons to *Pseudomonas* species using BLASTn against all 1071 full-length *Pseudomonas* genomes available from the *Pseudomonas* db v. 21.1 (2022-11-20) (Supplementary Table 4) ^66,77^. The best assignment was chosen according to BLASTn bitscores. We assigned *Z. tritici* reads of the *Z. tritici* amplicons on chromosomes 9 and 13, and of the ITS amplicon to a database of previously sequenced *Z. tritici* strains. For this, we used draft assemblies of previously collected 177 genomes from the Eschikon Field Station of the ETH Zurich, Switzerland ^37^, as well as genomes from the reference pangenome ^34^. The best assignment was chosen according to bitscores.

### Alignment, phylogenetic tree and network construction

We performed multiple sequence alignment using PASTA v.1.9.0 with the following MAFFT arguments --leavegappyregion --6merpair --maxiterate 0 --adjustdirection --reorder and FastTree model -gtr -gamma -fastest ^78^. We built phylogenetic trees of leaf samples using FastTree v.2.0.0 ^79^, and of the *Pseudomonas* mock samples using raxml-ng v. 1.2.0 with the options --all --model GTR+G --opt-model on --threads 4 --seed 2 --outgroup “Main ASV of *P. syringae* Leaf129” ^80^. To create unrooted phylogenetic networks for the *Z. tritici* amplicon 1 on chromosome 13, amplicon 2 on chromosome 9 and the ITS, the software SplitsTree v. 4.19.1 was used using uncorrected p distances ^81^.

### Statistics

Regression analyses were performed using the R function lm from the package stats v. 4.2.2 ^82^. Correlations were calculated using the R function cor.test from the package stats v. 4.2.2 ^82^. Shannon diversity was calculated using the function diversity from the R package vegan v. 2.6.4 ^83^. Permutations were calculated for 1000 iterations using the base R function sample with replace=TRUE ^71^. Subtrees were identified using the function subtrees from the R package ape v. 5.7.1 ^84^.

### Visualization

To visualize ASV abundances, the R package phyloseq v. 1.42.0 was used ^85^. Counts were normalized by sample using the phyloseq function transform_sample_counts(phyloseq_object, function (x) x / sum(x)). The heatmap was created using the R package pheatmap v. 1.0.12 ^86^. Table numbers were visualized with the method = "circle" of the R package corrplot v. 0.92 ^87^. Phylogenetic trees were visualized using the R package ggtree v. 3.6.2 ^88,89^. All other plots were created using the R package ggplot2 v. 3.4.2 ^90^. Organism and machine icons were created with BioRender.com.

## Author contributions

LS, MM and DC conceived the study, LS performed the research and analyzed the data, MM and DC supervised the work and acquired funding. LS and DC wrote the manuscript with input from MM.

## Data availability

Sequencing data generated for this work are available at the NCBI Sequence Read Archive (SRA). Analyzed datasets are provided as supplementary information.

## Acknowledgments

Data produced in this paper were generated in collaboration with the Genetic Diversity Centre (GDC), ETH Zurich. We thank Silvia Kobel and Aria Minder for the helpful discussions regarding the DNA extraction and PCR automations. We thank all the people who helped with the field sampling and processing, specifically Dominic Stalder, Nikhil Kumar Singh, Leen Abraham, Thomas Badet, Stéphanie Ruaud, Camille Kessler, Simone Fouché, Ursula Oggenfuss, Anna Spescha, Florence Gilliéron, Jana Schneider, Michael Brunner and Hansueli Zellweger. We thank Dominic Stalder and Sabina Tralamazza for all the helpful discussions on the analysis and Hanspeter Stalder for the input on PCR design and troubleshooting.

