## Supplementary Figures for "High-resolution Profiling of Bacterial and Fungal Communities using Pangenome-Informed Taxon-Specific Long-Read Amplicons"

Stalder et al.

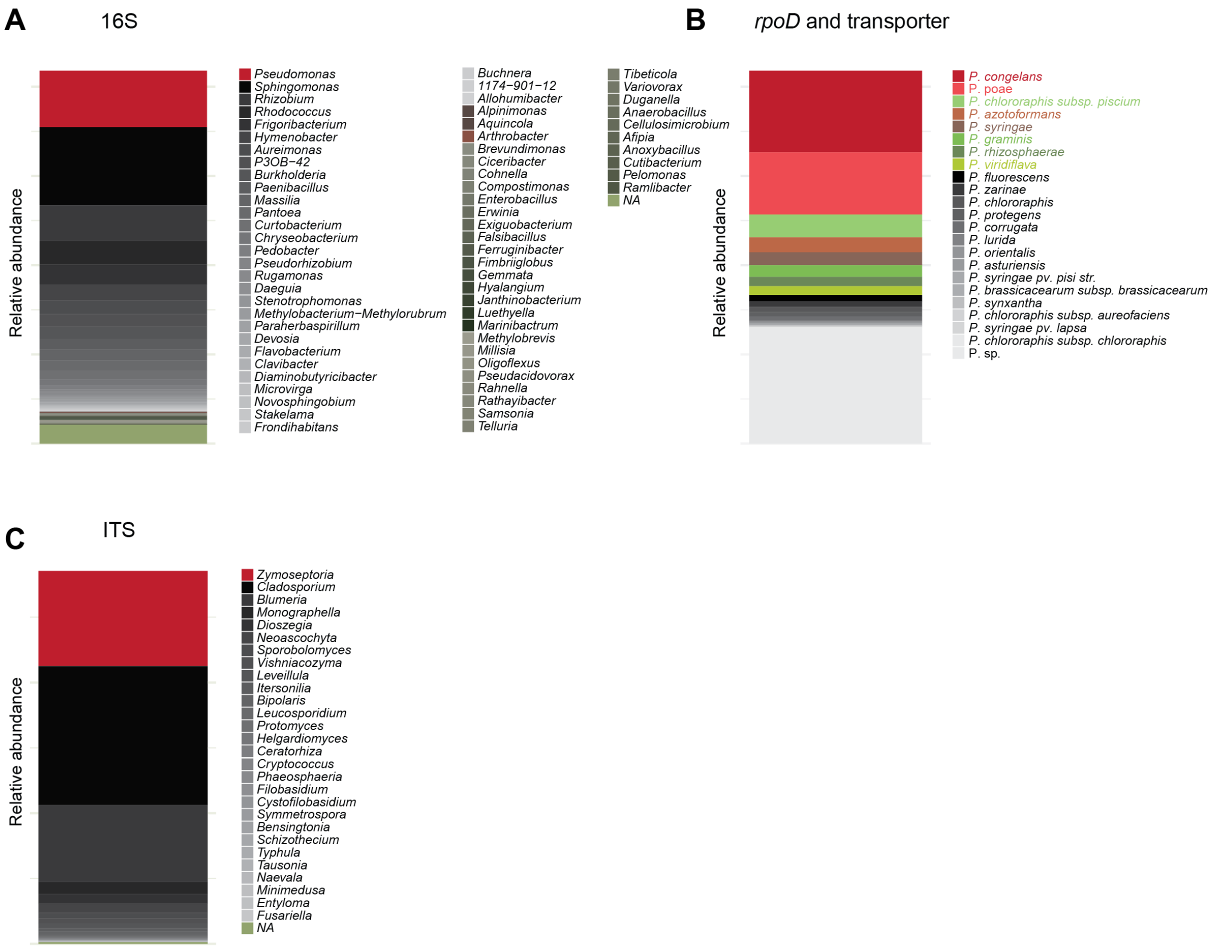


**Supplementary Figure S1: Relative abundances recovered by the 16S, *rpoD*, transporter and ITS amplicons.** (A) Relative abundances of bacterial genera on wheat leaves as determined by the full-length 16S amplicon. (B) Relative and mean abundances of *Pseudomonas* species on wheat leaves determined by the *Pseudomonas*-specific *rpoD* and transporter amplicons. (C) Relative abundances of fungal genera on wheat leaves as determined by the full-length ITS amplicon.

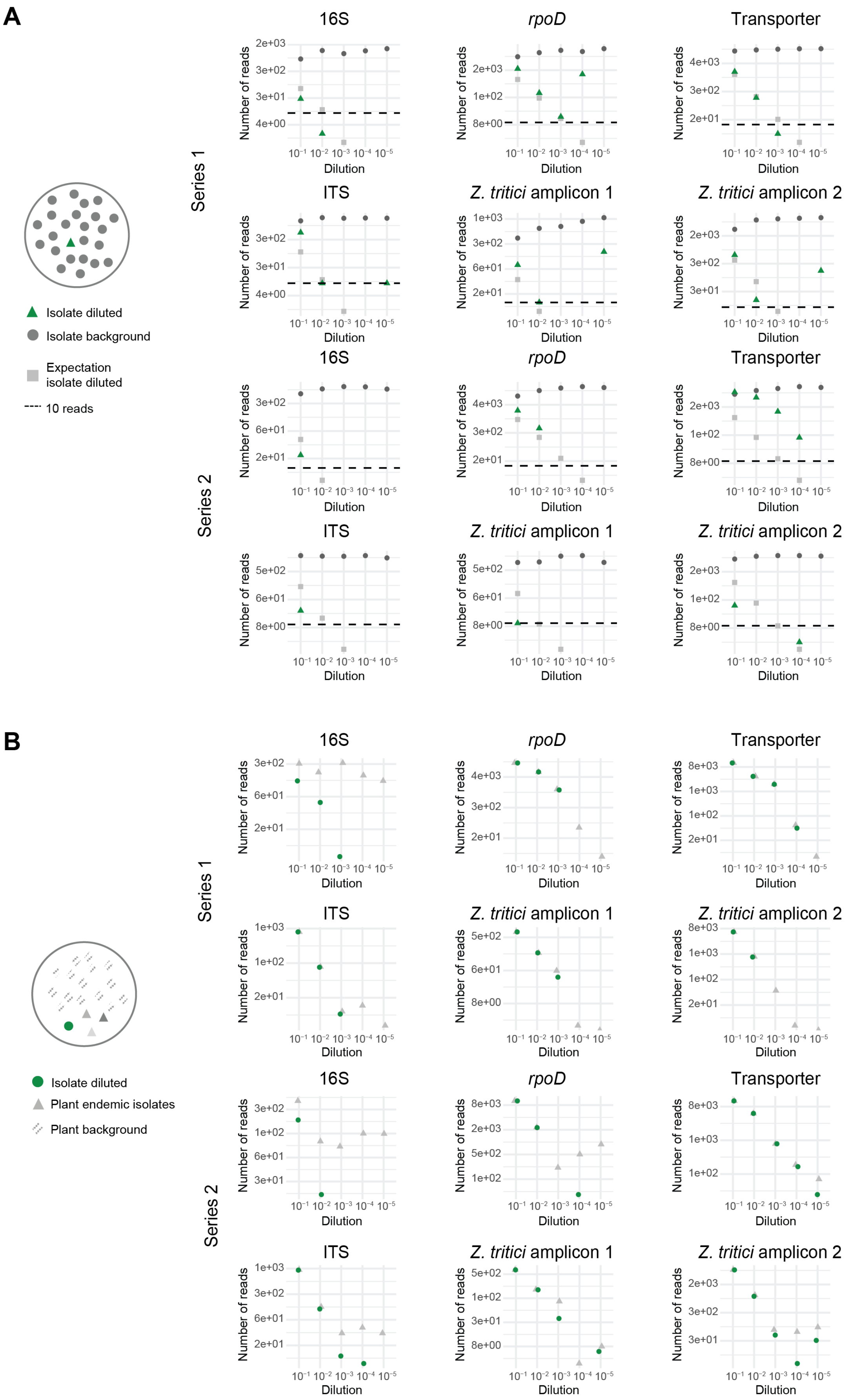

A
Series 1
Series 2
16S
rpoD
Transporter
ITS
Z. tritici amplicon 1
Z. tritici amplicon 2
Number of reads
Dilution
Isolate diluted
Isolate background
Expectation isolate diluted
10 reads
B
Series 1
Series 2
Isolate diluted
Plant endemic isolates
Plant background

**Supplementary Figure S2: Detection limit of taxon-specific amplicons using mock community analyses.**
(A) Series 1 - top: Serial dilution series of *P. syringae* Leaf129 (green) mixed into *P. thivervalensis* PITR2 (grey). Read numbers are shown for the full-length 16S, and the *Pseudomonas*-specific *rpoD* and transporter amplicons. The expected read numbers of the diluted *P. syringae* Leaf129 isolate are indicated in light grey and calculated from the background read numbers of *P. thivervalensis* PITR2. Series 1 - bottom: Serial dilution series of *Z. tritici* ST99CH 1E4 (green) in *Z. tritici* ST01IR 48b (grey) isolates. Read numbers are shown for the full-length ITS, and the *Z. tritici*-specific amplicons on chromosomes 9 and 13. The expected read numbers of the diluted *Z. tritici* ST99CH 1E4 isolate are indicated in light grey and calculated from the background read numbers of the *Z. tritici* ST01IR 48b isolate. Series 2: The same experiment as series 1 with diluted and background strains inverted.
(B) Series 1 - top: Read numbers of a serial dilution series of *P. thivervalensis* PITR2 (green) mixed into a wheat leaf DNA extract background for the full-length 16S amplicon and the *Pseudomonas*-specific *rpoD* and transporter amplicons. In grey, the total read count of the wheat-endemic bacterial isolates is indicated for the 16S amplicon, respectively the total read count of the wheat-endemic *Pseudomonas* for the *Pseudomonas*-specific *rpoD* and transporter amplicons. Series 1 - bottom: Read numbers of a serial dilution series of *Z. tritici* ST99CH 1E4 (green) mixed into a wheat leaf DNA extract background for the full-length ITS and the *Z. tritici*-specific amplicons. In grey, the total read count of the wheat-endemic fungal isolates is indicated for the ITS amplicon, respectively the total read count of the wheat-endemic *Z. tritici* for the *Z. tritici*-specific amplicons. Series 2: The same experiment as series 1, but performed with a different wheat leaf DNA extract background (see Methods). For all serial dilutions of (A) and (B), dilutions are performed with an initial DNA input of 7.5ng per isolate.

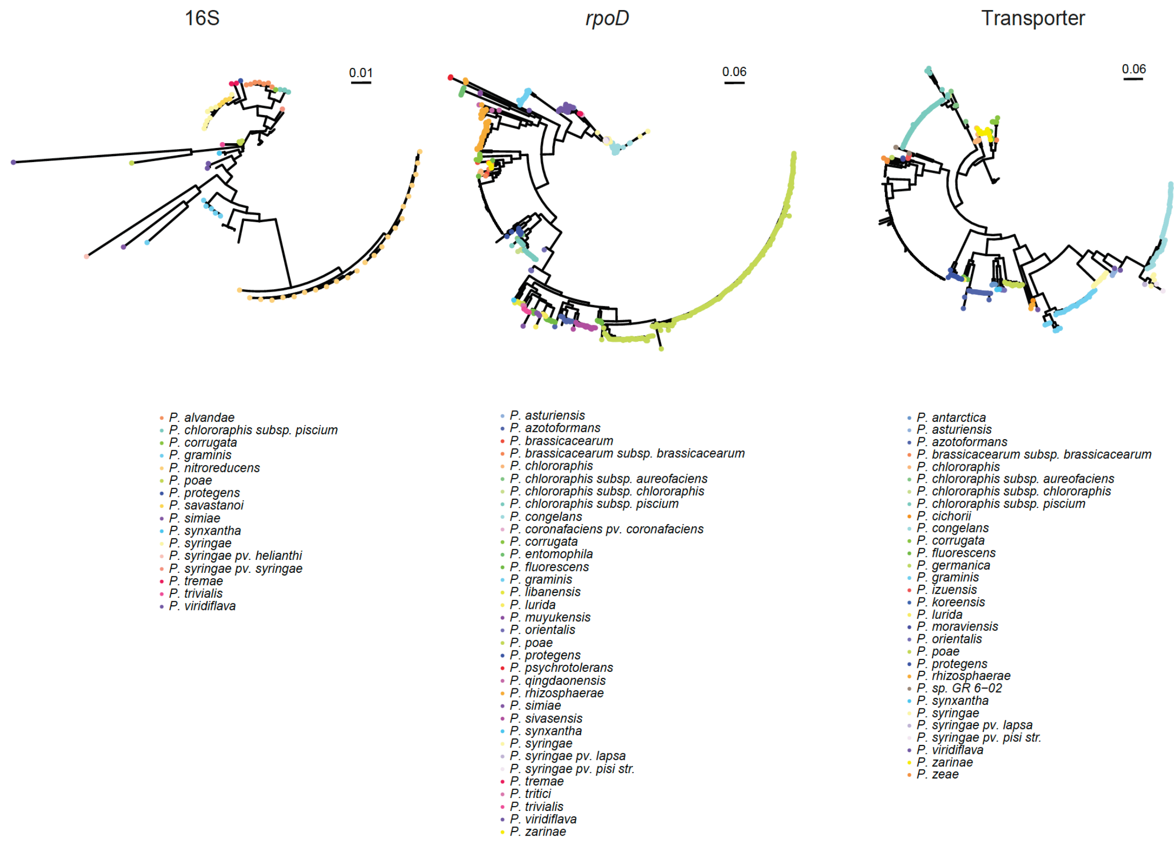


**Supplementary Figure S3: Resolution of taxon-specific amplicons in contrast to the full-length 16S.** Phylogenetic trees of *Pseudomonas* reads of the full-length 16S amplicon as well of the *Pseudomonas* specific *rpoD* and transporter amplicons. Reads are colored by *Pseudomonas* species.

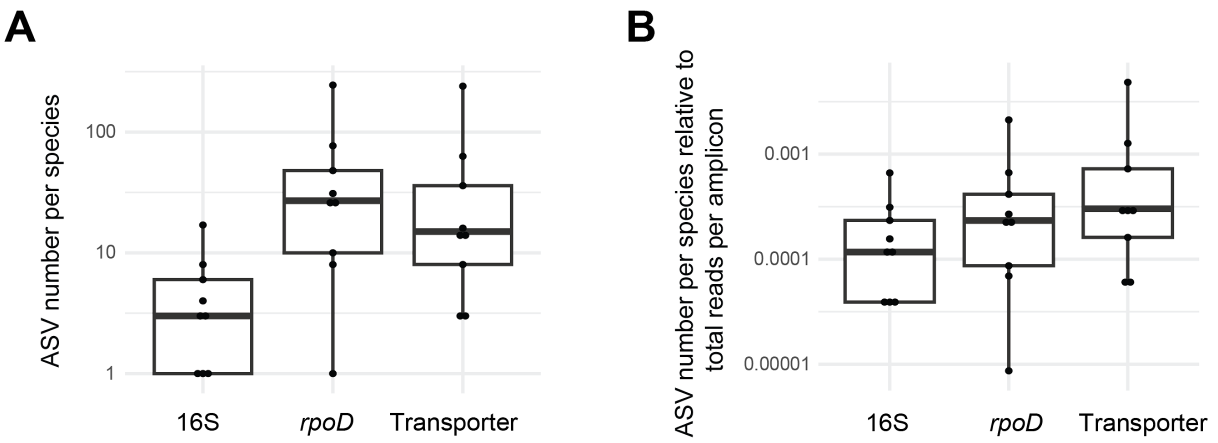


**Supplementary Figure S4: Number of amplicon sequence variants (ASVs) per species.** (A) Total number of ASVs per species for the full-length 16S amplicon, the *Pseudomonas*-specific *rpoD* and transporter amplicons. (B) Total number of ASVs per species divided by the total number of amplicon reads for the full-length 16S amplicon, the *Pseudomonas*-specific *rpoD* and transporter amplicons.

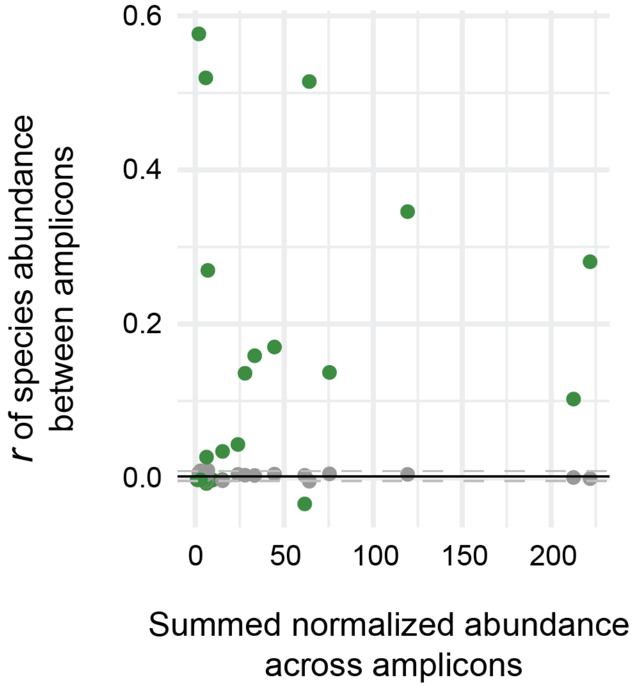


**Supplementary Figure S5: Species abundance correlation between the *rpoD* and transporter amplicons.** (A) Pearson correlation of *Pseudomonas* species abundance across samples between the *Pseudomonas*-specific *rpoD* and transporter amplicons. Correlation values for the species are compared to null expectations based on permutations (grey). The grey dotted line indicates the 95% confidence interval of the null expectation.

**A**

| | Genome size [Mbp] | Number of cores | Ribosomal diversity | Number of genomes | Number of OG | % genes in OG | % OG with all isolates | % OG with isolate-specific genes | Number of genomes |
|---|---|---|---|---|---|---|---|---|---|
| *Pseudomonas* | 6.5 | 1059 | 6.04 | 19 | 10026 | 96.6 | 16 | 0.5 | 19 |
| *Rhizobia* | 7.8 | 22 | 3.15 | 84 | 19600 | 98.4 | 5.2 | 0.1 | 49 |
| *Streptomyces* | 8.5 | 2784 | 1.29 | 94 | 13148 | 97.3 | 11 | 0.2 | 94 |
| *Z. tritici* | 39.7 | 6389 | 0.01 | 19 | 13975 | 99.5 | 65.8 | 0.1 | 19 |
| *A. fumigatus* | 29.4 | 5132 | 0.67 | 253 | 11411 | 99.9 | 69.4 | 0 | 252 |

**B**

| | Genome size [Mbp] | Number of cores | Ribosomal diversity | Number of genomes | Number of OG | % genes in OG | % OG with all isolates | % OG with isolate-specific genes | Number of genomes |
|---|---|---|---|---|---|---|---|---|---|
| *Pseudomonas* | 6.5 | 1059 | 5.1 | 19 | 10026 | 96.6 | 16 | 0.5 | 19 |
| *Rhizobia* | 7.8 | 22 | 12.65 | 19 | 11516 | 95.7 | 16.7 | 0.5 | 19 |
| *Streptomyces* | 8.5 | 515 | 6.04 | 19 | 9643 | 95.3 | 18.5 | 0.5 | 19 |
| *Z. tritici* | 39.7 | 6389 | 0.01 | 19 | 13975 | 99.5 | 65.8 | 0.1 | 19 |
| *A. fumigatus* | 29.4 | 5107 | 7.24 | 19 | 10023 | 99.8 | 84.8 | 0 | 19 |

**Supplementary Figure S6: Pangenome characteristics of the bacterial groups *Pseudomonas*, *Rhizobia*, and *Streptomyces*, as well as the fungal species *Z .tritici* and *A. fumigatus*.**
(A) Pangenome characteristics of *Rhizobia*, *Streptomyces*, *Pseudomonas*, *Z. tritici* and *A. fumigatus*. The pangenome of *Rhizobia* is based on the genome set of Yang et al. (2020), the pangenome of *Streptomyces* on the genome set of Kieper et al. 2023 and the pangenome of *A. fumigatus* on the genome set of Barber et al. 2021.
For each pangenome, the average genome size in Mbp, the number of candidate core regions and the ribosomal diversity, as defined by the length-normalized nucleotide diversity of the 16S or ITS regions, is indicated. In addition, genes were grouped into orthogroups based on protein homology for each pangenome. The total number of orthogroups, the percentage of genes included in orthogroups, the percentage of orthogroups that contain all isolates, and the percentage of orthogroups that contain isolate-specific genes are specifically indicated. (B) Same as (A), but pangenomes were calculated based on a subset of 19 representative genomes per taxon group for better comparability.

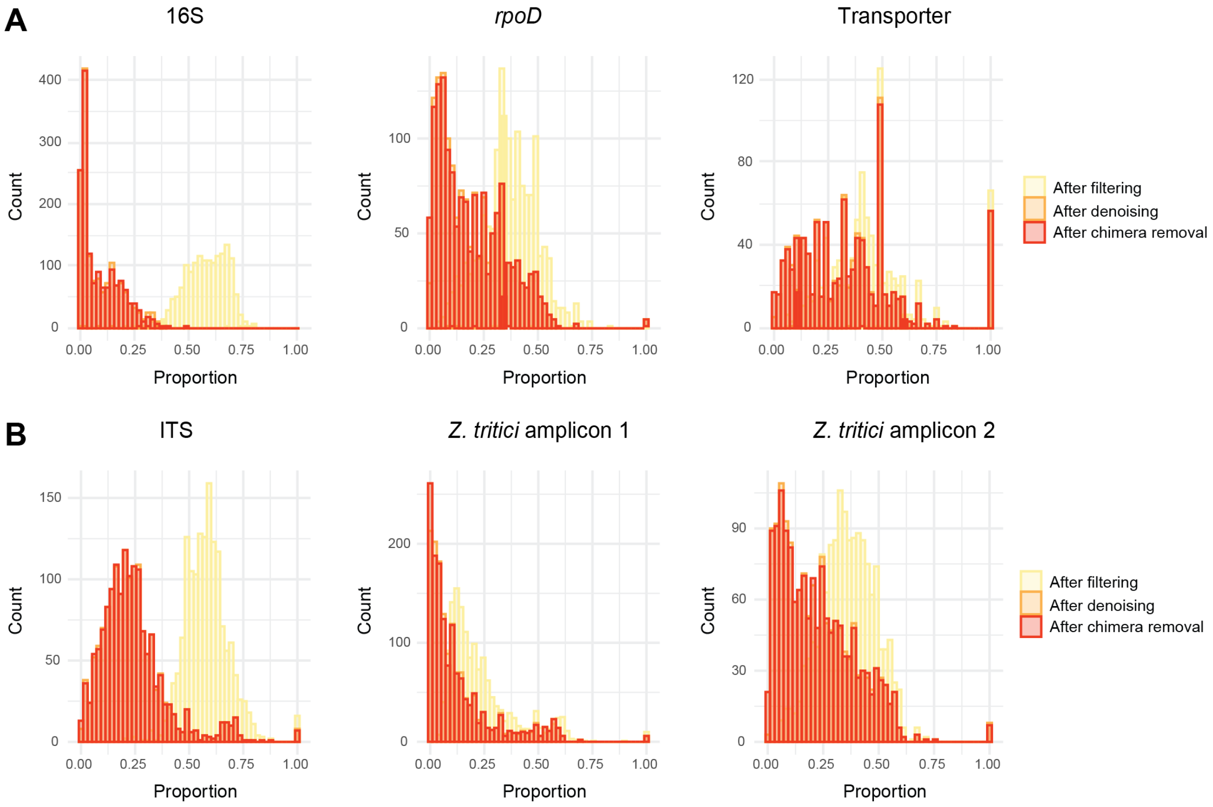


**Supplementary Figure S7: Read tracking through the dada2 pre-processing pipeline.** Proportion of reads retained after dada2 filtering, after dada2 denoising and after dada2 chimera removal for (A) the full-length 16S amplicon, and the *Pseudomonas* specific *rpoD* and transporter amplicons and for (B) the full-length ITS amplicon and *Z. tritici*-specific amplicons 1 and 2.
